# Population-wide variation in connectopic organization of cerebral language hubs

**DOI:** 10.64898/2026.03.12.711274

**Authors:** Jitse S. Amelink, Carmen Ramoser, Sourena Soheili-Nezhad, Dick Schijven, Meng-Yun Wang, 23andMe Research Team, Simon E. Fisher, Christian F. Beckmann, Koen V. Haak, Clyde Francks

## Abstract

The uniquely human capacity for language is supported by a distributed brain network with cerebral cortical hubs in the left inferior frontal gyrus (LIFG) and left superior temporal sulcus (LSTS). Individual differences in the graded spatial organization of function across these regions may provide sensitive phenotypes for behavioural and genetic association analyses, to better understand the biological underpinnings of language. In 41,437 UK Biobank participants we applied connectopic mapping to the LIFG and LSTS. This approach maps spatial changes across a region in terms of connectivity with the rest of the brain, based on resting state functional imaging. On average, we found that Brodmann Area (BA) 44, BA 45 and the posterior LSTS were strongly connected to the superior and inferior parietal lobe and the dorsal attention and frontoparietal networks, whereas BA 47 and the medial LSTS were strongly connected to superior frontal areas BA 8B and BA 9, and the default mode network more broadly. However, there was extensive population-wide variation in these graded connectopic patterns, which significantly mediated the effects of polygenic scores for reading ability and educational attainment on performance on a vocabulary task. Furthermore we identified 3 genomic loci displaying multivariate association patterns with these connectopic maps, one of which coincides with a long non-coding RNA, *LINC01165*, with potential relevance in human evolution. Overall, connectopic mapping produced fewer significant genetic association findings and lower heritability (range from 0.3 to 9.4%) than previous parcel-based approaches for mapping language network connectivity. These observations suggest higher genetic complexity and/or prominent roles for environmental or random developmental effects on these connectopic maps.

## Introduction

The human capacity for language is unique in its complexity among forms of communication in the animal kingdom^1–3^. The functional and structural organization of various brain regions is adapted to support language^2,4,5^. The involvement of the inferior frontal gyrus and superior temporal sulcus in lanugage was already reported in the 19th century^6^. Modern experimental work using functional magnetic resonance imaging (fMRI) has characterised these areas in greater detail through diverse analytic and experimental strategies in relation to language tasks^7–10^. Task-based individual maps have been shown to be highly similar to the resting state network connectivity of these regions^11–14^. In fact spatial changes in seed-based resting state functional connectivity patterns can be indicative of boundaries between functionally distinct brain areas^15–17^, showing the relevance of local-global connectivity patterns for parsing brain organization.

Population scale mapping of functional connectivity in previous neuroimaging genome-wide association studies (GWAS) investigated genetic contributions to global resting state properties of the language network^18–22^ and the brain more broadly^23^. These studies identified associations of functional connectivity with various genomic loci, and with overall polygenic dispositions to language-related traits such as the reading disorder dyslexia. However, the imaging anaysis methods in these studies did not capture spatially graded changes in local-global connectivity patterns, but rather relied on atlas-based parcellation. To the extent that connectivity gradients exist within key hubs of the brain’s language network, then mapping their average forms and individual differences in large-scale population data may help to reveal further aspects of the neurobiology and genetic underpinnings of brain language organization.

Connectopic mapping is a computational approach specifically designed to map functional heterogeneity within a region, and to capture this at an individual level. The method disentangles overlapping organizational modes of functional connectivity within given a region of interest, by mapping within such a region the connectivity profile for each voxel with the rest of the brain, and then deriving latent patterns to describe the spatial connectivity profiles. Here, we applied a toolbox called CONGRADS^17,24–26^ to imaging data from 41,437 participants of UK-Biobank to generate connectopic maps within the key language-related areas in terms of their respective connectivity modes with the rest of the brain, and thereby also measured inter-individual differences in this organization.

We aimed to answer the following questions: First, what is the average connectopic organization of the LIFG and LSTS in the population? Second, how do these cortical language hubs vary in their connectopic organization across individuals in large-scale population data? Third, to what extent are the inter-individual differences heritable, might they mediate associations between vocabulary size (i.e. the only directly language-related measure available in UK Biobank from the majority of participants who have brain image data) and polygenic scores for language-related traits such as dyslexia, and which specific genomic loci are involved? Fourth, given that language is a uniquely human characteristic, what is the evolutionary history of the implicated genomic loci?

Specifically, we measured individual differences in the first and second connectopic maps for each region (LIFG and LSTS). In total we computed 12 imaging derived phenotypes (IDPs) from these connectopic maps using independent component analysis^27^ and used these to investigate population-wide variation in local-global patterns. Eleven of these IDPs showed significant SNP-based heritability, and we tested the associations of these 11 IDPs with polygenic scores and picture vocabulary task performance, as well as investigating their genetic architecture in multivariate GWAS (mvGWAS), followed by an investigation of the evolutionary history of the implicated loci. See Figure 1 for a study overview.

**Figure 1:**
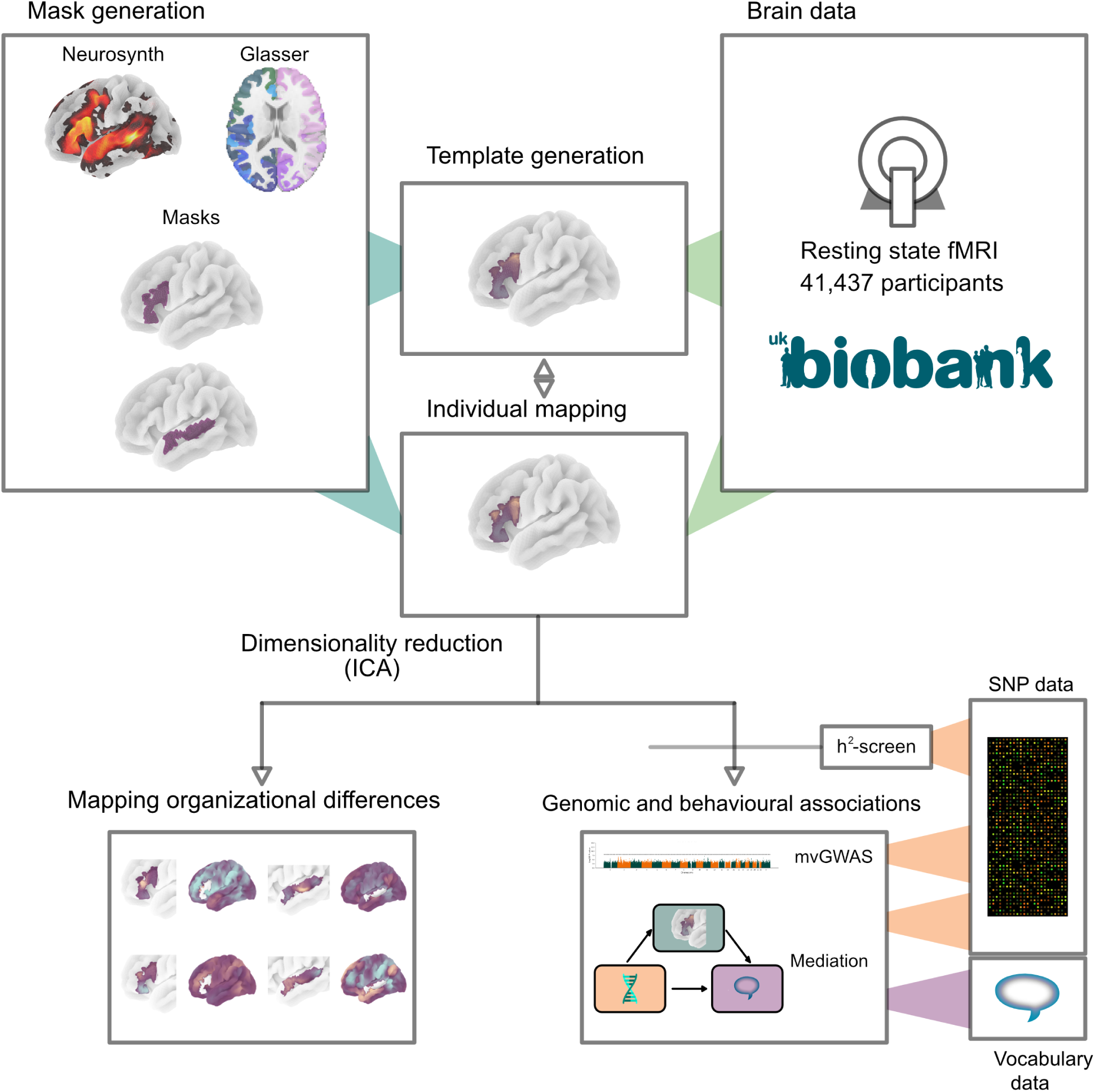
Study overview. Study overview. We derived language-related region-of-interest (ROI) masks based on Neurosynth results and applied connectopic mapping (CONGRADS) to derive a template based on 998 randomly selected subjects. We then derived and aligned individual connectopic maps based on this pipeline for 41,437 participants. We used 11 heritable imaging-derived phenotypes (IDPs) to test genomic and behavioural associations using polygenic scores, vocabulary performance and genome-wide associations.

## Results

### Estimating connectopic maps in UK Biobank

The first step for our approach was to identify cortical regions-of-interest (ROIs) involved in language processing, to be used for deriving connectopic maps. We downloaded the meta-analytic results related to the keyword “language” from Neurosynth (v.7)^8^, and parcellated these results using the multimodal Glasser atlas^15^. We identified 11 parcels that were highly associated with language (*P <* 1 ∗ 10*^−^*^8^), and that could be combined into two distinct ROIs: one in and near the left inferior frontal gyrus (LIFG, atlas parcels 74-6, 79 and 81-2, all in cortex region 21) and the other in and near the left superior temporal sulcus (LSTS, atlas parcels 125, 128-130, all in cortex region 11, see Supplementary Figure 1 for a step-by-step breakdown). The extension beyond the narrow classical language areas (e.g. Brodmann Area (BA) 44 and 45) is in line with findings from subject-specific mapping of language sensitivity^10^.

We then proceeded to compute the connectopic maps using CONGRADS^24^ on a population-scale. There were 41,437 participants of UK-Biobank that passed genetic and imaging sample quality control (see Methods), and for which we computed all connectopic maps. In brief, CONGRADS derives overlapping organizational modes by comparing the within-ROI time courses with connectivity fingerprints of the rest of the brain, and modeling the low-dimensional organizational modes using manifold learning^24^. As overlapping modes of organization are present in the cortex^17,24,28^, we derived the first and second connectopic map within each ROI (connectopic map - CMAP) and its projected connectivity pattern to the rest of the brain (projected map - PMAP). Individual CMAPs for each subject were aligned to the overall CMAP template using spatial correlation. After that,the CMAPs were used for IDP derivation using MELODIC^27^, and the PMAPs were used to inspect associated whole-brain connectivity patterns. We investigated both the population average CMAP and PMAP, by annotating the distribution of the PMAP across the 7 main brain networks classified by Yeo and colleagues^29^. As the Yeo annotation does not list a specific language network, for analysis of each ROI (LIFG or LSTS) we added the other ROI as an extra category.

Connectopic maps provide insights into distinct connectivity patterns within both ROIs. The first connectopic map (henceforth “G1” for brevity) in the LIFG ROI runs from the superior part of the ROI (inferior frontal junction) to the inferior part (the lateral part of BA 47) (Figure 2, Supplementary Table 1). The dorsal positively loaded part of the connectopic map shows strongest connectivity to the superior and inferior parietal areas, corresponding to the dorsal attention network, whereas the inferior part of the ROI shows strongest average connectivity to two superior frontal areas (lateral part of BA 8B and medial part of BA 9), both in the default mode network (Figure 2, Supplementary Table 1). The second connectopic map (henceforth “G2”) of the LIFG ROI runs from the middle part of the ROI (BA45) to both ends of the ROI (Supplementary Figure 3) and also displays strong connectivity to both superior and inferior parietal areas and the dorsal attention network (Supplementary Figure 2, Supplementary Table 1).

**Figure 2:**
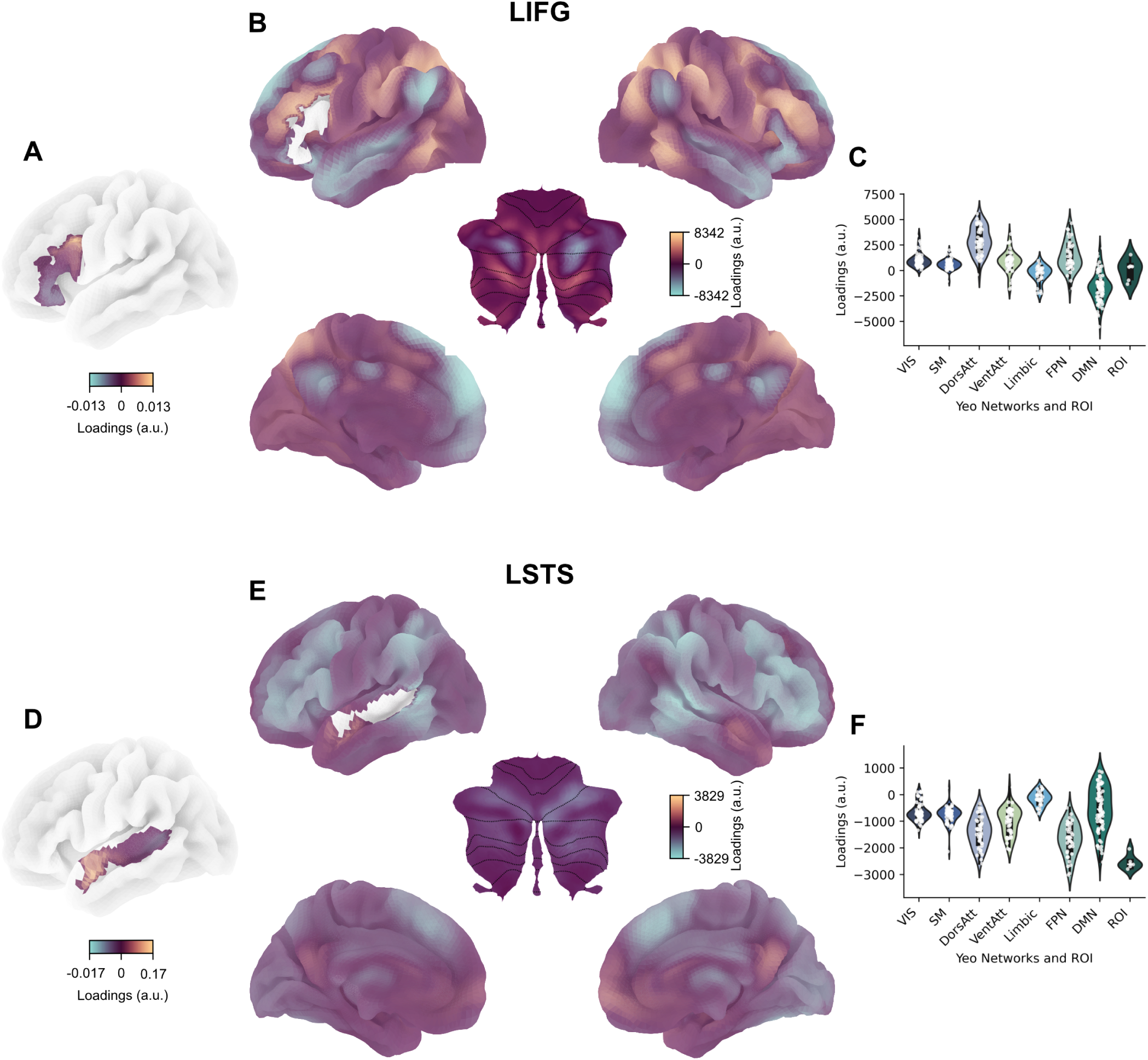
Population-average connectopic maps and their connectivity patterns. **A + D.** The first average connectopic map (CMAP) within each ROI. **B + E.** Average connectivity patterns for the first CMAP of each ROI to the rest of the brain (projected map - PMAP). **C + F.** Network distributions of the first PMAPs based on the Yeo 7 networks and the Glasser atlas. **A-F.** Note that the directions of loadings are arbitrary, but that the sign corresponds between the CMAPs and the PMAPs, meaning that green regions in A+D correspond to green regions in B + E. Abbreviations: LIFG – Left inferior frontal gyrus. LSTS – left superior temporal sulcus. VIS - visual network. SM - somatomotor network. DorsAtt - dorsal attention network. VentAtt - ventral attention network. Limbic - limbic system. FPN - frontoparietal network. DMN - default mode network. ROI - region of interest (corresponding to the LSTS ROI for subplot C and to the LIFG ROI for subplot F). a.u. - arbitrary units.).

For the temporal lobe, the first connectopic map of the LSTS runs anterior to posterior, with the posterior part displaying strong connectivity to the LIFG ROI and inferior parietal areas (Figure 2, Supplementary Table 1), which are part of the frontoparietal network. The anterior part displays weak connectivity to the posterior and anterior cingulate. The second connectopic map of the LSTS runs again from the middle of the ROI to the anterior and posterior parts. It is most strongly connected to superior frontal areas (lateral part of BA 8B and medial part of BA 9) and anterior and posterior cingulate, which are part of the default mode network (Supplementary Figure 2, Supplementary Table 1).

### Generating imaging derived phenotypes from connectopic maps

Next we reduced the dimensionality of the CMAPs using FSL’s MELODIC^27^ in order to derive IDPs for genomic and behavioural follow-up analyses. Three components was the optimal model order across connectopic maps (Supplementary Figure 3). Spatial and subject loading based correlations showed distinct clustering patterns (Supplementary Figure 4-5), with the subject loadings clustering primarily across ROIs, such that temporal and frontal functional brain organizations are clearly related.

To investigate population-wide variation in functional brain organization of the brain language areas, we investigated the ends of the distribution of the independent components. We averaged CMAPs and PMAPs of the bottom 50, middle 50 and top 50 individual participants for each component (Figure 3, Supplementary Figure 6, Supplementary Table 2). The middle 50 are fairly representative of the population average. For most of the components we observe one end that shows strongly distinct PMAP connectivity patterns, while the other end of the component does not show a distinguishable connectivity pattern. Two components (Figure 3, G1 LIFG IC 2 and 3) display a very sharp boundary towards the posterior part of the inferior frontal gyrus in the upper 50 subjects that indicates a smaller functional language area and which is associated with a less pronounced connectivity pattern (Figure 3). Other components capture the decoupling of BA44 and BA45 (Figure 3, G1 LIFG IC 1, G2 LIFG IC 1). In the temporal ROI, the first component (Figure 3, G1 LSTS IC1) captures a functional shift of the anterior STS upwards. The second component in this ROI (Figure 3, G1 LSTS IC2) signifies a distinct shift in connectivity patterns of the anterior and posterior STS, with the lower end showing integration between the anterior STS and the default mode network, whereas in the upper end the anterior STS integrates with visual and motor networks.

**Figure 3:**
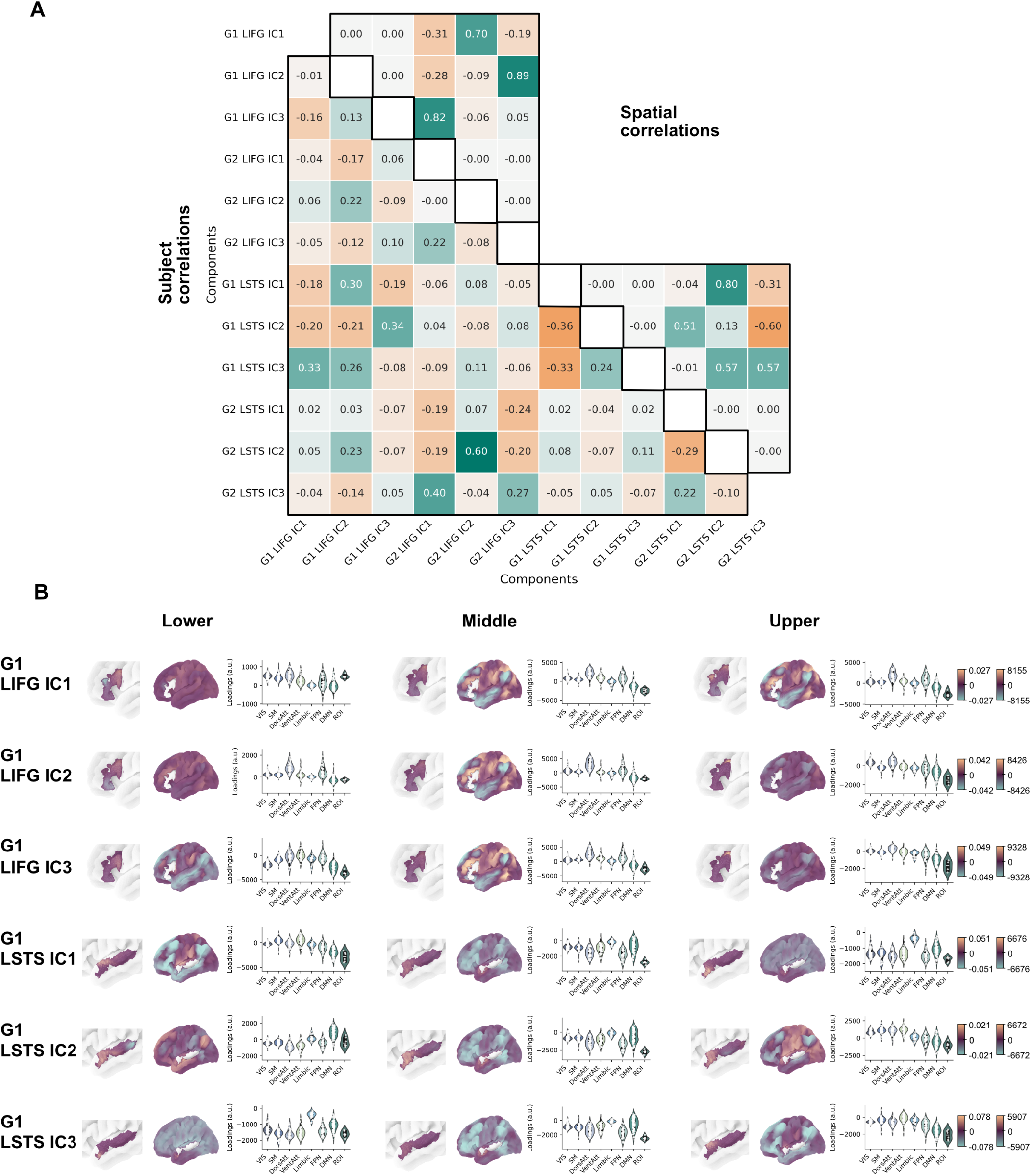
Phenotypic correlations and population variation in brain organization. **A.** Subject-based (bottom) and spatial correlations (top) of the independent components for CMAP 1 and 2 (G1 and G2). **B.** Population-based variation of CMAP and PMAP differences. The average CMAPs and PMAPs are shown for connectopic map 1 (G1) for the bottom 50, middle 50 and upper 50 individual participants on the IC loadings. Network preferences per IC by Yeo 7 networks. Abbreviations: LIFG – Left inferior frontal gyrus. LSTS - left superior temporal sulcus. VIS - visual network. SM - somatomotor network. DorsAtt - dorsal attention network. VentAtt - ventral attention network. Limbic - limbic system. FPN - frontoparietal network. DMN - default mode network. ROI - region of interest (corresponding to the LSTS ROI for the top three rows of figure B and to the LIFG ROI for the bottom three rows of figure B). a.u. - arbitrary units.

### Behavioural and polygenic associations of connectopic maps

We then estimated the SNP-based heritability of the 12 IDPs derived using MELODIC^27^ (3 independent components for 2 connectopic maps in 2 brain regions) using GCTA^30^ and estimated the inverse-variance weighted mean heritability over all of these IDPs. SNP-based heritability estimates were in the range of 0.3-9.4% (Supplementary Table 3), with an inverse variance-weighted mean of 3.3%. 11 out of 12 IDPs had a nominally significant SNP-heritability and were used for further analysis.

We next investigated whether functional brain organization mediates the effect of genetic dispositions for language-relevant traits and neurodevelopmental conditions on vocabulary performance, using the same approach as previous work^22^. We calculated polygenic scores (PGS) in the same cohort of 41,437 UK Biobank participants. Traits for which we calculated PGS included reading ability^31^, left-handedness^32^, educational attainment^33^, dyslexia^34,35^, autism^36^, attention deficit/hyperactivity disorder (ADHD)^37^ and schizophrenia^38^, based on summary statistics available from prior large-scale GWAS efforts.

Before performing mediation analyses, we tested all associations using a generalized linear model that included sex, age, genetic and imaging covariates (Methods). After correcting for 95 tests, we identified 20 significant associations with the IDPs: 11 that passed Bonferroni correction, another 9 that passed FDR-correction (*q <* 0.05). 6 of these associations were found with the PGS for reading. Another 2 associations were identified with the PGS for educational attainment. Finally one association was found with the PGS for dyslexia and one IDP (G2 LSTS IC 2) (Figure 4, Supplementary Table 4). 5 out of 11 IDPs were also associated with vocabulary performance (N=20,976, Figure 4, Supplementary Table 5). Higher PGS for performance or vocabulary scores were associated with more pronounced differences in connectivity patterns, with a greater degree of similarity and integration. We additionally observed that all polygenic scores, except for left-handedness, showed associations with vocabulary level (N=20,976, Figure 4, Supplementary Table 6).

**Figure 4:**
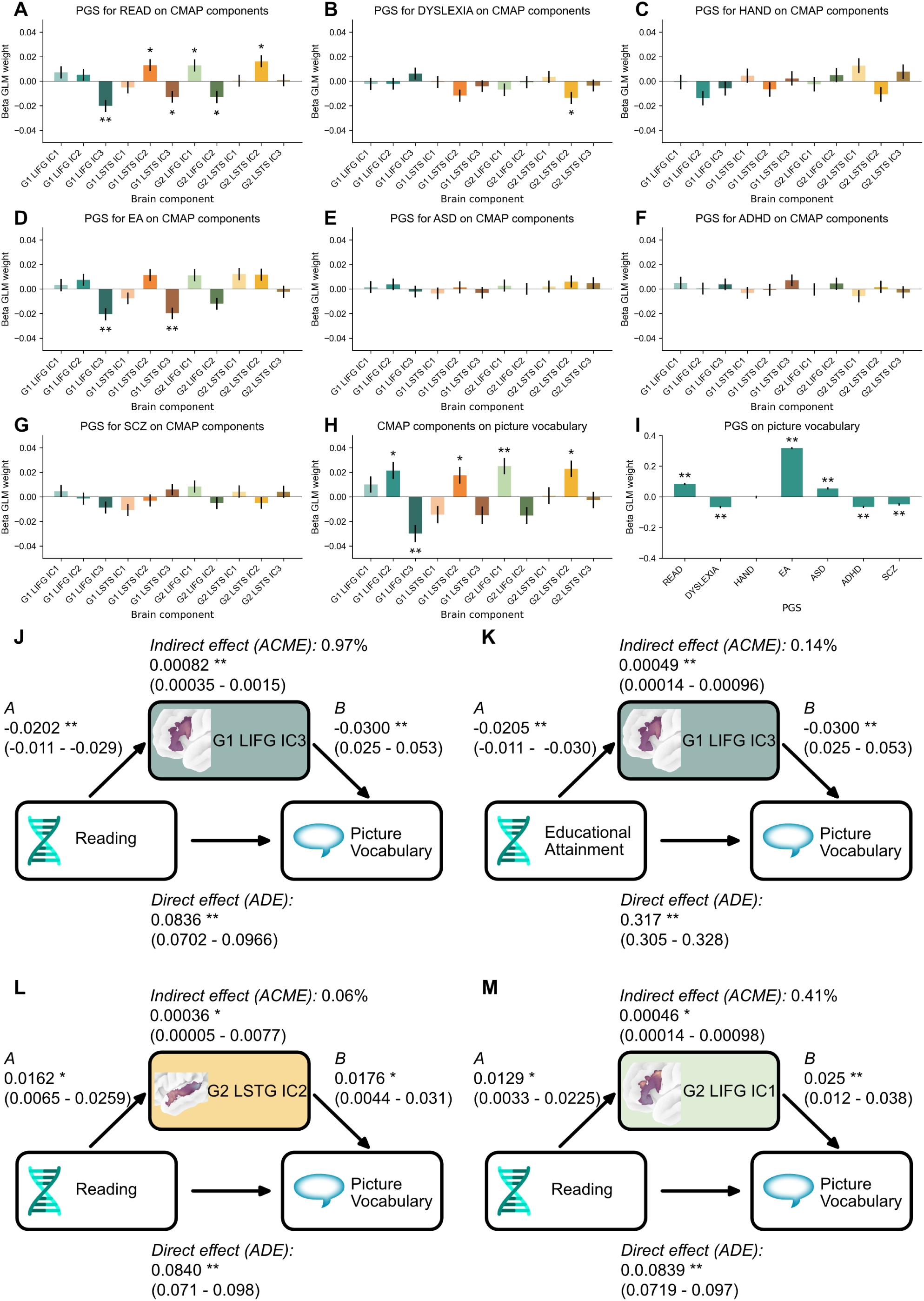
Polygenic and behavioural associations of connectopic maps. **A-G.** Polygenic score associations for 7 language-related traits with 11 heritable IDPs (N=41,437). **H.** Brain IDP associations with vocabulary performance (N=20,976). **I.** Polygenic score associations with vocabulary performance (N=20,976). **J-M.** Mediation analyses for interaction effects that were significant in the original pairwise associations (N=20,976). ** indicates significance after Bonferroni correction for 95 independent tests in **A-I** and 24 tests in **J-M**, * indicates False Discovery Rate-corrected significance (q < 0.05) for respectively 95 and 24 tests.

In total this resulted in 6 mediation models for testing. We applied the mediation package in R (v.4.4.0) to 20,976 participants with 1,000 permutations. 4 out of 6 of the tested mediation models showed a significant indirect mediation path (Figure 4, Supplementary Table 7). The two sets of analyses with the most significant mediation effects involved G1 LIFG IC 3 and polygenic scores for reading and educational attainment respectively.

### Genetic correlates of connectopic maps

We then conducted a multivariate GWAS using REGENIE v.3.6.0^39^ on the 11 heritable IDPs. We identified three genome-wide significant loci (*P <* 5 ∗ 10*^−^*^8^, Figure 5), one of which has not been previously identified in a brain imaging GWAS. All three loci showed multivariate association patterns across IDPs and were not driven by one IDP only (Figure 5). We did not identify any significant enrichment when testing for gene-set enrichment with MAGMA for 12,477 gene sets as implemented in FUMA (Methods) (Supplementary Tables 8-12, Supplementary Figures 7-10).

**Figure 5:**
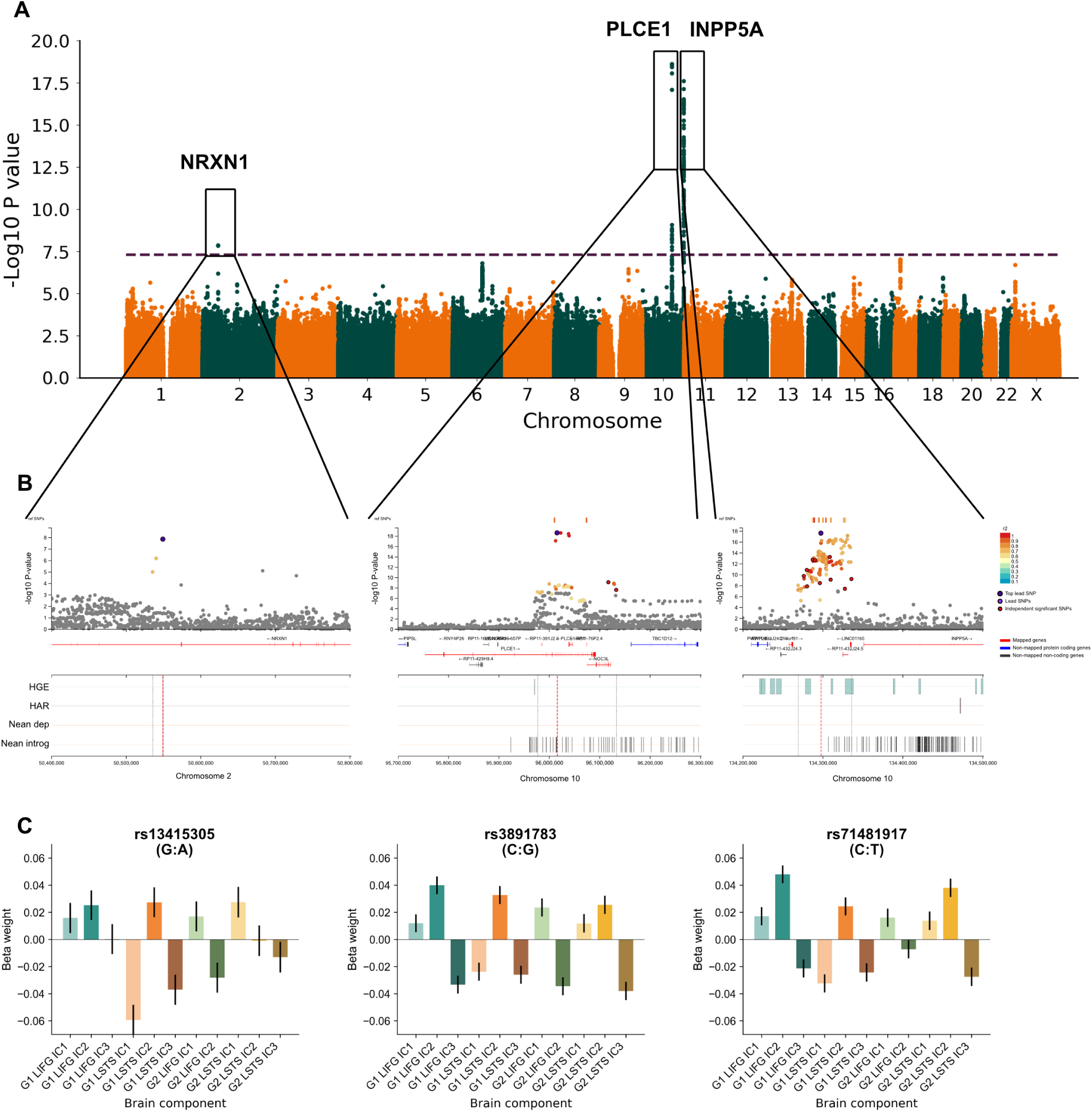
Molecular correlates and evolutionary signatures of genomic loci associated with functional brain organization. **A.** Multivariate GWAS (mvGWAS) plot of 11 heritable IDPs. Dashed line indicates genome-wide significance (*P* = 5 ∗ 10*^−^*^8^). X-axis indicates location on the genome, Y-axis indicates association significance by -log10 P-value. **B.** Locuszoom plots of three lead SNPs with overlapping evolutionary annotations for fetal brain human gained enhancers (HGE), human accelerated regions (HARs), archaic deserts (Nean depleted) and Neandertal introgressed (Nean introgressed) regions. **C.** Beta weights for lead SNPs with all 11 IDPs, showing multivariate association profiles.

To examine the evolutionary and functional characteristics of the genetic loci identified by the mvGWAS, we looked up the lead SNPs in the Human Genome Dating Atlas^40^ and GTEx^41^ for expression quantitative trait loci (eQTLs). In addition, we also conducted an overlap analysis with four evolutionarily relevant annotations of the human genome: Neandertal-introgressed fragments^42^, archaic deserts (regions significantly depleted for Neandertal introgressed alleles)^43^, fetal brain human-gained enhancers (HGEs)^44^ and human accelerated regions (HARs)^45–48^. These annotations span an array of different evolutionary timelines, from HGEs describing genomic changes that date back as much as approximately 30Mya, and the Neandertal introgressed regions and archaic deserts dating back 50-60,000 years ago, with HARs describing genomic loci that are conserved across vertebrates, but diverged in the human lineage after the human-chimpanzee split (approximately 6 Mya) a broader range in between. Note that an overlap analysis is a hypothesis-generating, rather than statistical confirmatory test.

The lead SNP rs13415305 on chromosome 2 is intronic to *NRXN1* and not previously identified in a brain imaging GWAS according to the GWAS Catalog^49^. *NRXN1* encodes a cell surface protein that is part of the neurexin family and is involved in cell-adhesion and neurotransmitter release through Ca^2+^ binding. No eQTLs have been reported for this SNP and the locus does not show overlap with genomic regions of interest defined by our evolutionary annotations. Rare deleterious mutations in *NRXN1* are associated with a broad variable neurodevelopmental syndrome including language deficits, schizophrenia and autism^50,51^. The derived A allele for this polymorphism dates back approximately 30,000 years, therefore relatively recently in human evolution.

The locus on chromosome 10 near *PLCE1* and *NOC3L* is well-documented for its association with functional language connectivity^20–23^. *PLCE1* encodes a phospholipase that generates two second messengers: inositol 1,4,5- triphosphate (IP3) and diacylglycerol (DAG). *NOC3L* (previously labelled *C10orf117*) encodes a protein that is similar to nucleolar complex-associated 3 (NOC3) and is involved in RNA-binding. The associated genomic locus contains Neandertal introgressed alleles (Supplementary Table 13). The lead SNP rs3891783 is annotated as an eQTL for *NOC3L* in brain tissue (lowest *P* = 3 ∗ 10*^−^*^6^ for the amygdala), and the lead SNP is an eQTL for both *NOC3L* and *PLCE1* in the tibial nerve (*P* = 6.2 ∗ 10*^−^*^10^ and *P* = 6.3 ∗ 10*^−^*^7^). The derived C allele dates back roughly 1,38 Mya, indicating a relatively deep presence for this polymorphism in the human lineage.

The final locus on chromosome 10 near *INPP5A* and *LINC01165* is also well-described for association with functional language connectivity^20–23^. *INPP5A* is a membrane-associated phosphatase that generates IP3, the same second messenger also derived from *PLCE1*. *LINC01165* is a long intergenic non-coding RNA, of which the function is currently unclear. The broader locus contains 3 fetal brain HGE elements that span around 25% of the locus and also contains multiple Neandertal introgressed alleles (Supplementary Table 13). The lead SNP rs71481917 is an eQTL in brain tissue for *LINC01165* (*P* = 2.7 ∗ 10^11^) and *INPP5A* (*P* = 1.2 ∗ 10*^−^*^9^) and the derived T allele (reference: C) dates back roughly 500,000 years.

## Discussion

Understanding variation in brain functional organization is one of the great challenges in the neurobiology of language and neuroscience more broadly. In this study we characterized individual differences and functional heterogeneity within key language-related cortical hubs in the left hemisphere, by applying connectopic mapping to these regions. From the connectivity patterns emerged two clusters: BA 44, BA 45 and the posterior STS were strongly connected to the superior and inferior parietal lobe and the dorsal attention and frontoparietal networks. BA 47 and the medial STS on the other hand were strongly connected to superior frontal areas BA 8B and BA9, although we observed population-wide variation in multiple components that describe variation in this organization. In our genomic and behavioural analyses we found that several components partly mediated the effects of polygenic scores for reading ability and educational attainment on vocabulary level. Furthermore we identified 3 independent genomic loci, that all displayed multivariate association patterns with the IDPs, of which one locus encompassed fetal brain human gained enhancer elements.

An advantage of using gradual spatial changes in connectivity, rather than hard parcellations, was that it helped to tease apart the organization and population variation within core language-related regions. The first connectopic map (G1) of the LIFG identified a gradient from BA 44 through to BA 47. BA 45 was more distinct in the second connectopic map (G2), but also showed more similarity in connectivity profile with BA44 in some of the participants, and more with BA47 in others, with the population-average leaning towards similarity with BA 44. In addition, we could identify sharp boundaries that indicated a smaller BA44 and BA45, which we found to be associated with lower polygenic scores for reading and educational attainment and lower vocabulary levels.

For the LSTS we could tease apart the posterior (G1) and medial parts (G2). The posterior part showed strong connectivity to the LIFG ROI, which is in line with previous work that suggests that the posterior part is an important language hub^52^. The integration of both BA44 and BA45 and the posterior STS with superior and inferior parietal areas is largely aligned with the Memory Unification Control model^25,53,54^ and fits the meta-analytic finding that these parts are more strongly activated by syntactic processing^5,55,56^. The integration of BA47 and the medial STS with parts of the default mode network fits the notion of more semantic processing happening there^5,55,56^.

The connectivity patterns identified in this study showed subtle but significant associations with genetic dispositions affecting language-related traits (reading, educational attainment and dyslexia) and to the most directly language-related cognitive test available in UK Biobank, vocabulary performance. The polygenic score for reading ability, as in our previous work based on integrated brain structure-function analysis^22^, was the most significantly associated with brain variability in both LIFG and LSTS. Concerning the direction of the effects, polygenic dispositions to higher performance / reduced risk of disorder was always associated with a more pronounced connectivity pattern in the brain, rather than a flatter connectopic map. This is in part due to averaging: on the other tail there are opposing effects, i.e. a positive and negative loading that cancel each other out, but our findings can also indicate weaker spatial connectivity changes with higher genetic risk of language-related disorders. Stronger brain integration into distinct networks might therefore be beneficial for language and other cognitive abilities. Overall, the association profiles for vocabulary scores and PGS for reading ability, and to a lesser extent educational attainment, were highly similar. Apart from a few associations for PGS for dyslexia, all other PGS showed no associations. It should be noted that all of these associations are subtle, and differences in significance should be interpreted with caution.

Investigating the histories of the genetic loci implicated in GWAS studies of language-related traits may shed light on the evolution of language^3,57^. From an evolutionary perspective, the *LINC01165* locus identified in our GWAS is especially interesting. Non-coding RNAs may have been critical in human brain evolution^58,59^, through their effects on regulation of gene expression. Lower expression in macaques of homologues of human long non-coding RNAs in the brain compared to protein-coding genes^60^ also suggests a potential role for long non-coding RNAs in brain evolution^61^. *LINC01165* lies entirely in a fetal brain HGE region. Although lead SNP rs71481917 is not in a fetal brain HGE region, it is an eQTL for *LINC01165* throughout the brain according to the latest GTEx-release. However, apart from its association with resting-state fMRI^23^, little is known about the functions of this gene. It may be worth investigating more systematically the potential contributions of long non-coding RNAs to the evolution of the language-ready brain, by studying their associations with brain structural/functional variability in modern-day humans, and also through functional genomic analysis of cell lines and brain organoids.

Heritability estimates of connectopic IDPs and the number of genome-wide significant loci from GWAS were lower in this study compared to previous functional connectivity work based on parcellation of core language regions^21–23^. However, the association strength of the IDPs in the present study with vocabulary performance is highly similar to previously derived brain structure-function language components, which were not derived based on within-region functional variability^22^. These findings imply that finegrained functional organization within cortical regions might be less genetically constrained than at a larger scale, with greater contributions from environmental factors or chance processes during development, but with similar mediating effects between polygenic dispositions and cognitive performance^62,63^. Additionally, parcel-based approaches may be more affected by heritable anatomical rather than functional differences. Another recent study of functional gradients also showed limited GWAS-based discovery^64^, albeit not using the local-global approach employed in CONGRADS. An additional factor may be that transmodal association areas have been reported to have lower heritability of structure-function coupling than unimodal sensory areas^65^. All these factors may have contributed to the overall lower genetic signal observed in the present work.

Our study has a number of limitations. First, UK Biobank is a relatively healthy cohort that is not completely representative of the whole British population^66,67^. Resting-state fMRI metrics are generally less heritable than structural measures in UK-Biobank data^68^, although this varies per analysis strategy^68^ and may also be related to a relatively short scanning time in UK Biobank, which might not be optimal for brain-wide estimation^69^. In addition, in line with previous connectopic mapping work^24^, we excluded subjects with a low similarity to the study template, because these maps likely reflect data artefacts and not underlying biology^70^ and because highly dissimilar maps are incomparable at population scale. It is nonetheless possible that some of these maps do reflect true biological variation.

In conclusion, we have identified average patterns and population-wide variation in the functional organization of the LIFG and LSTS. The local-global variation allowed us to identify integration with areas in the parietal part of the frontoparietal and dorsal attention networks for the posterior LSTS and the posterior LIFG, and to the default mode network for the medial LSTS and anterior part of the LIFG. We observed that one of the mapped genes that is significantly associated with this variability, *LINC01165*, is in a fetal brain HGE region, suggesting a potential role in brain evolution. Overall we observed lower SNP-based heritability estimates than in previous studies that used different imaging phenotypes^21,22,68^, suggesting that connectopic maps of cerebral cortical language hubs are affected to a greater extent by environmental and/or chance developmental effects in brain development.. All of this supports a probabilistic rather than deterministic understanding of how the human genome affects brain language organization.

## Methods

### Participants

As described previously^21^, imaging and genomic data were obtained from the UK Biobank^71^ as part of research application 16066 from primary applicant Clyde Francks. UK Biobank received ethical approval from the National Research Ethics Service Committee North West-Haydock (reference 11/NW/0382), and all of their procedures were performed in accordance with the World Medical Association guidelines. Informed consent was obtained for all participants^72^. Analyses were conducted on 41,437 participants that remained after quality control (QC) of genotype and imaging data (see below).

### Imaging data

Brain imaging data were collected for UK Biobank as described previously^73,74^. Identical scanners and software platforms were used for data collection at different sites (Siemens 3T Skyra; software platform VD13). For collection of resting state fMRI data, participants were instructed to lie still and relaxed with their eyes fixed on a crosshair for a duration of 6 minutes. In that timeframe 490 datapoints were collected using a multiband 8 gradient echo EPI sequence with a flip angle of 52 degrees, resulting in a TR of 0.735 s with a resolution of 2.4x2.4x2.4mm^3^ and field-of-view of 88x88x64 voxels. Our study made use of pre-processed image data generated by an image-processing pipeline developed and run on behalf of UK Biobank^73^.

### Genetic data

Genome-wide genotype data (UK Biobank data category 263) were obtained by the UK Biobank using two different genotyping arrays (for full details see^71^). Imputed array-based genotype data contained over 90 million SNPs and short insertion-deletions with their coordinates reported in human reference genome assembly GRCh37 (hg19). In downstream analyses we used both the non-imputed and imputed array-based genotype data in different steps (below).

### Sample-level quality control and ancestry estimation

First, participants with available imaging data (N = 63,620, release March 2025) were extracted from the full UK Biobank imputed genotype dataset^71^. Subject level QC parameters were then applied to the 61,910 individuals for whom both neuroimaging and genotype data were available. This involved excluding individuals with a mismatch of their self-reported (UKB data field 31) and genetically inferred sex (UKB data field 22001), as well as individuals with putative aneuploidies (UKB data field 22019), or individuals who were determined as outliers based on heterozygosity (PC corrected heterozygosity *>* 0.1903) or genotype missingness rate (missing rate *>* 0.05) (UKB data field 22027). Next, we identified pairs of individuals with a kinship coefficient *>* 0.0442 (UKB data field 2202111), and excluded one individual from each pair, prioritizing the exclusion of individuals related to a larger number of other individuals to maximize the overall sample. 59,537 individuals remained after these filtering steps.

For ancestry estimation, we used Neural Admixture^75^, a neural network autoencoder, to calculate admixture estimates for every individual. First, we harmonised genotypes of filtered individuals with those of the 1000 Genomes Phase 3^76^. We used those genotypes and superpopulation labels (AFR, EAS, EUR, SAS, AMR) of 1000 Genomes individuals for supervised training of the neural net. The trained net was used to infer admixture of filtered individuals. For a balance between maximal sample size and homogeneity, a threshold of 97% EUR ancestry was chosen in addition to a PC1 cutoff defined by 1000 Genomes EUR. 56,226 individuals remained after genetic sample QC.

In phenotype sample-level QC, we first excluded participants with imaging data labeled as unusable by UK Biobank QC. Second, participants were removed based on outliers (here defined as 3 standard deviations from the mean) and missingness in at least one of the following metrics: discrepancy between resting state fMRI brain image and T1 structural brain image (UK Biobank field 25739), inverted temporal signal-to-noise ratio in preprocessed and artefact-cleaned preprocessed resting state fMRI (data fields 25743 and 25744), scanner X, Y and Z brain position (fields 25756, 25757 and 25758) or with poor alignment from the CMAP template (*r >* 0.3, see Imaging data preprocessing and phenotype derivation). We excluded 9,679 participants based on these QC steps, as well as an additional 4,980 based on missing imaging derived phenotypes (IDPs) due to poor spatial alignment to one or more of the 4 CMAP templates. In total 41,437 participants without any missing values were included in all main analyses.

### Imaging data pre-processing and phenotype derivation

Preprocessing was conducted by the UK Biobank and consisted of motion correction using MCFlirt^77^, intensity normalization, high-pass filtering to remove temporal drift (sigma=50.0s), unwarping using fieldmaps and gradient distortion correction. Structured scanner and movement artefacts were removed using ICA-FIX^27,78,79^. Preprocessed data were registered to a common reference template in order to make analyses comparable (the 6th generation nonlinear MNI152 space, http://www.bic.mni.mcgill.ca/ServicesAtlases/ICBM152NLin6) and smoothed with a Gaussian filter (*σ* = 2.5) using FSL^80^.

Region-of-interest (ROI) masks were derived by parcellating the meta-analytic map for language from Neurosynth^8^ with the Glasser atlas^15^. Parcels that were most strongly associated with language (*P <* 1 ∗ 10*^−^*^8^) were combined into one cluster in the left inferior frontal gyrus (LIFG) and one cluster in the left superior temporal sulcus (LSTS), and eroded around the edges to make the ROI smoother, to avoid having part of the ROI outside the brain. We used these two ROI masks for deriving connectopic maps.

Connectopic mapping is a method to tease apart modes of connectivity change within a ROI^24^. The analysis procedure consists of two main steps: computing a similarity matrix between the correlations of the time courses of the ROI and those of the rest of the brain using *η*^2^, and derive the Laplacian Eigenmaps. The resulting within-ROI maps (“CMAPS”) are then regressed back on to the resting-state fMRI data from the rest of the brain (“PMAPS”) to gain insight into what connectivity patterns underpin the modes of connectivity change in the ROI.

In order to align each individual’s CMAPs and PMAPs to the same template, we first computed reference templates from the averaged similarity matrix from resting state data from 998 randomly selected UK-Biobank subjects. We then derived individual CMAPs and PMAPs and aligned them to this CMAP template using spatial cross-correlation^80^ between the individual CMAPs and template CMAPs and flipped both CMAPS and PMAPS in case of negative correlations, since the sign of the connectopic map is arbitrary. We used a cut-off of *r >* 0.3 for inclusion, as low spatial similarity makes the maps incomparable (Supplementary Figure 11).

In order to derive imaging phenotypes from these maps, we derived independent component analysis (ICA) of the CMAPs using MELODIC from FSL v. 6.0.3 on one-fourth of the participants that were each time randomly selected. To derive these components a 4D matrix was entered into MELODIC, with each subject 3D CMAP concatenated along the fourth dimension. The optimal order was estimated to be 3 components based on principal component analysis (Supplementary Figure 3), and was therefore applied in all analyses. We computed IDPs using 3 independent components for 2 connectopic maps in 2 brain regions, yielding 12 IDPs, which were compared for SNP-based heritability.

For post-hoc functional annotation and interpretation, population average PMAPs were parcellated using the Glasser atlas^15^ and annotated using Yeo 7 Networks^29^.

### Genetic variant-level QC

Three different genetic datasets were prepared, as needed for four different analysis processes:

1. Array-based genotype data were filtered, maintaining variants with linkage disequilibrium (LD) ≤ 0.9, minor allele frequency (MAF) ≥ 0.01, Hardy-Weinberg Equilibrium test *P*-value ≥ 1 × 10*^−^*^15^ (see^39^), and genotype missingness ≤ 0.01 for REGENIE step 1 (below).
2. Imputed genotype data were filtered, maintaining bi-allelic variants with an imputation quality ≥ 0.7, Hardy-Weinberg Equilibrium test *P*-value ≥ 1 × 10*^−^*^7^ and genotype missingness ≥ 0.05 for association testing in REGENIE step 2.
3. For genetic relationship matrices, SNPs were only used if they were bi-allelic, had a genotype missingness rate ≤ 0.02, a Hardy Weinberg Equilibrium *P*-value ≥ 1 × 10*^−^*^6^, an imputation INFO score ≥ 0.9, a MAF ≥ 0.01, and a MAF difference ≤ 0.2 between the imaging subset and the whole UK Biobank.

### Heritability analysis

Genetic relationship matrices (GRMs) were computed for the study sample using GCTA v. 1.93.0beta^30^. In addition to the previous sample-level QC, individuals with a genotyping rate ≤ 0.98 and one random individual per pair with a kinship coefficient ≥ 0.025 derived from the GRM were excluded from heritability analysis. The SNP-based heritability of all 12 IDPs was estimated using genome-based restricted maximum likelihood (GREML) in GCTA v. 1.93.0beta^30^. Category-level heritability was computed using the inverse-variance weighted mean. We used all nominally significant (*P <* 0.05) heritable phenotypes for downstream analyses.

### Mediation analysis

We tested (1) the association between polygenic scores for seven language-related traits and picture vocabulary scores (UK Biobank field 26302-2-0), (2) the association between polygenic scores and our brain language IDPs, (3) the association between the IDPs and picture vocabulary performance, before (4) testing to what extent the IDPs mediate the association of polygenic scores with picture vocabulary scores.

The picture vocabulary test in UK Biobank is a computer adaptive test to measure a participant’s vocabulary level, adapted from the NIH Toolbox^81^ and recalibrated for use in a UK cohort^82^. All of the words within the test are rated with a difficulty score from -11 to 11. In each round, the participant is shown a word on a screen and has to choose the correct picture for that word. If the picture is chosen correctly, the participant will get a more difficult word next round, if not, the next word will be easier. Picture vocabulary is a reliable indicator of linguistic knowledge^83^. The picture vocabulary test is the only test of a linguistic cognitive function available in UK Biobank for more than 5000 participants.

Polygenic scores in UK Biobank individuals were calculated based on SNP-wise effect sizes from previous GWAS studies: for reading abilities (quantitatively assessed in up to 33,959 participants from the GenLang consortium)^31^, dyslexia (51,800 cases and 1,087,070 controls) from 23andMe Research Institute^34^ and left-handedness (UK Biobank participants without imaging data: 33,704 cases and 272,673 controls)^32^, educational attainment (765,283 individuals)^33^, autism spectrum disorder (18,381 cases, 27,969 controls)^36^, attention deficit hyperactivity disorder (38,691 cases, 186,843 controls))^37^ and schizophrenia (76,755 cases, 243,649 controls)^38^.

We derived polygenic scores with PRS-CS^84^ and used a generalized linear model and mediation analysis (using the package mediation^85,86^) in R (v 4.40) to test these questions. PRS-CS, a Bayesian regression framework to infer posterior effect sizes of autosomal SNPs based on genome-wide association summary statistics, was applied using default parameters and a recommended global shrinkage parameter phi = 0.01, combined with LD information from the 1000 Genomes Project phase 3 European-descent reference panel. PRS-CS performs to a similar level as other polygenic scoring methods, with noticeably better out-of-sample prediction than a clumping and thresholding approach^87,88^. Covariates included in the model were genetic sex, age, age^2^, age × sex, the first 10 genetic principal components that capture genome-wide ancestral diversity, genotype array (binary variable) and various scanner-related quality measures (Supplementary Table 14). Associations between PGS and IDPs were tested in 41,437 participants, whereas the other analyses were conducted in all 20,976 participants with all IDPs and picture vocabulary scores available.

### Genome-wide association testing

REGENIE v.3.6.0 was used^39^ for genome-wide association testing. In brief, REGENIE is a two-step machine learning method that fits a whole genome regression model and uses a block-based approach for computational efficiency. In REGENIE step 1, array-based genotype data were used to estimate the polygenic signal in blocks across the genome with a two-level ridge regression cross-validation approach. The estimated predictors were combined into a single predictor, which was then decomposed into 23 per-chromosome predictors using a leave one chromosome out (LOCO) approach, using a block size of 1000, 4 threads and low-memory flag. These LOCO polygenic predictors were used as covariates in both common and rare variant association testing in REGENIE step 2. Phenotypes were transformed to a normal distribution in both REGENIE step 1 and 2. Covariates for both steps included genetic sex, age, age2^^^, agesex, the first 10 genetic principal components that capture genome-wide ancestral diversity^71^, genotype array (binary variable) and various scanner-related quality measures (scanner X, Y and Z-position, inverted temporal signal to noise ratio and mean displacement as an indication of head motion, amount of T1 warping to standard space, discrepancy between resting state fMRI brain image and T1).

In REGENIE step 2, we performed separate single-variant association analyses with common (imputed) variants separately for the broad and narrow brain language components using a Firth regression framework. GWAS were run on our local compute cluster using a block size of 500, 4 threads and a low-memory flag. We ran the multivariate GWAS using the multiphen-flag.

Genome-wide significant variants, Bonferroni-adjusted for genome-wide signficance (i.e. threshold *P <* 5 ∗ 10*^−^*^8^), were annotated using the online FUMA platform (version 1.5.2)^89^. MAGMA (version 1.08)^90^ was used to calculate gene-based P-values and to investigate potential gene sets of interest^91,92^ and to map the expression of associated genes in a tissue-specific^41^ and time-specific^93^ fashion. Gene sets smaller than 10 were excluded from the analysis.

### Evolutionary overlap analysis

We conducted an overlap analysis with the following evolutionary annotations. An overlap analysis is a hypothesis-generating analysis, rather than a confirmatory analysis.

1. Fetal brain human-gained enhancers (HGEs)^44^. HGEs were previously defined across three consecutive developmental periods in human fetal cortical tissue^44,94^ compared to mice and marmosets.
2. Neandertal introgressed alleles^42^. We used a previously derived annotation that includes both Neandertal-introgressed SNPs, and those in perfect linkage disequilibrium (*r*^2^ = 1), reflecting a relatively recent (35,000-50,000 years ago) evolutionary event (i.e., Neandertal-Homo sapiens admixture).
3. Archaic deserts^43^. We used previously defined archaic deserts that represent genomic regions with significant depletion of Neandertal sequence (average introgression percent < 10^−3.5^) across Europeans, East Asians, South Asians, and Melanesians. From the six identified desert regions spanning 85.3 Mb, we eliminated 128 Neandertal introgressed SNPs (constituting approximately 10*^−^*^4^ of the total desert length) along with haplotypes in substantial LD (*r*^2^ *>* 0.6).
4. Human accelerated regions (HARs)^45–48^ are regions in the genome that are conserved among vertebrates, but show accelerated evolution on the human lineage.

### Data and code availability

The primary data used in this study are from UK Biobank. These data can be provided by UK Biobank pending scientific review and a completed material transfer agreement. Requests for the data should be submitted to UK Biobank: https://www.ukbiobank.ac.uk. Specific UK Biobank data field codes are given in Materials and Methods. Dyslexia GWAS summary statistics must be obtained by request to 23andMe Research Institute, via https://research.23andme.com/dataset-access/.

Other publicly available data sources and applications are cited in Materials and Methods. We have made our mvGWAS summary statistics available online within the GWAS catalog: https://ebi.ac.uk/gwas/. This study used openly available software and codes, specifically GCTA (https://cnsgenomics.com/software/gcta/ #GREML), FUMA (https://fuma.ctglab.nl/), MAGMA (https://ctg.cncr.nl/software/magma, also implemented in FUMA), PRS-CS (https://github.com/getian107/PRScs), REGENIE (https://rgcgithub.github.io/regenie/install/), Neural Admixture (https://github.com/AI-sandbox/neural-admixture), CONGRADS (https://github.com/koenhaak/congrads) and FSL (https://fsl.fmrib.ox.ac.uk/fsl/docs/). Custom code for this study is available from https://github.com/jsamelink/congrads_genetics. All other data needed to evaluate the conclusions in the paper are present in the paper and/or the Supplementary Materials.

## Acknowledgements

This research was funded by the Max Planck Society (Germany). Brain imaging, genomic and cognitive data were obtained from UK Biobank as part of research application 16066 from primary applicant Clyde Francks. Our study partly made use of pre-processed image data generated by an image-processing pipeline developed and run on behalf of UK Biobank. The funders had no role in study design, data collection and analysis, and the decision to publish or preparation of the manuscript. The authors would like to thank the participants of UK Biobank. The authors would like to thank the research participants and employees of 23andMe Research Institute for making this work possible. The authors thank Gökberk Alagöz and Else Eising for their input on the evolutionary analyses.

## Author Contributions

Conceptualization - J.S.A, C.F.B, K.V.H., S.E.F., C.F. ; Methodology - J.S.A, G.A., C.R., C.F.B., K.V.H., D.S.; Software - J.S.A., S.S-N., G.A., A.L., D.S.; Formal analysis - J.S.A., C.R; Data curation - J.S.A, C.R., M-Y. W., 23andMe; Writing - original draft - J.S.A.; Writing - review & editing - C.R., G.A, D.S., K.V.H., S.E.F, C.F.B., C.F ; Visualization - J.S.A.; Project administration - C.F.; Resources - S.E.F, C.F.; Funding acquisition - C.F., S.E.F.; Supervision - K.V.H, S.E.F., C.F.B., C.F.

## Author information

### 23andMe Research Team

Adam Auton, Alan Kwong, Anjali J. Shastri, Barry Hicks, Catherine H. Weldon, David A. Hinds, Emily DelloRusso, Emily M. Rios, Joyce Y. Tung, Kahsaia de Brito, Katelyn Kukar Bond, Keng-Han Lin, Matthew H. McIntyre, Matthew J. Kmiecik, Qiaojuan Jane Su, Robert K. Bell, Sayantan Das, Shubham Saini, Stella Aslibekyan, Vinh Tran, Wanwan Xu, Alisa P. Lehman, Noura S. Abul-Husn, R. Ryanne Wu, Rebecca M. K. Berns, Ruth I. Tennen, Stacey B. Detweiler, Aditya Ambati, Anna Guan, Bertram L. Koelsch, Chris German, Éadaoin Harney, Ethan M. Jewett, G. David Poznik, James R. Ashenhurst, Jingran Wen, Peter R. Wilton, Steven J. Micheletti, and William A. Freyman.

## Disclosures

C.F.B. is director and shareholder of SBGneuro Ltd. The other authors have no competing interests to declare.

