## Supplementary Figures for "Population-wide variation in connectopic organization of cerebral language hubs"

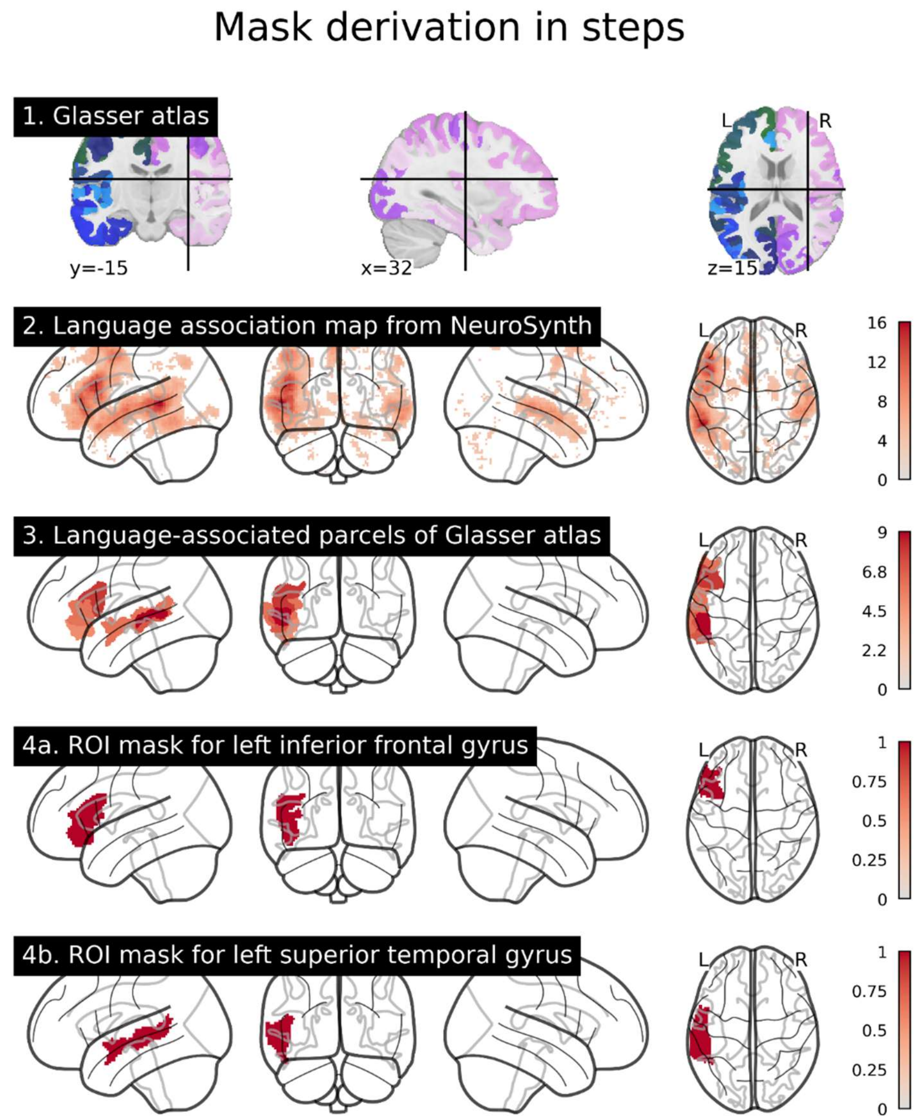


**Supplementary Figure 1.** Mask derivation broken down in steps. We combine the multimodal Glasser atlas (1) with the language association map from Neurosynth (2) to find which brain areas are most strongly associated with language task performance (3). These are then combined in two region-of-interest (ROI) masks (4a and 4b) for further analysis.

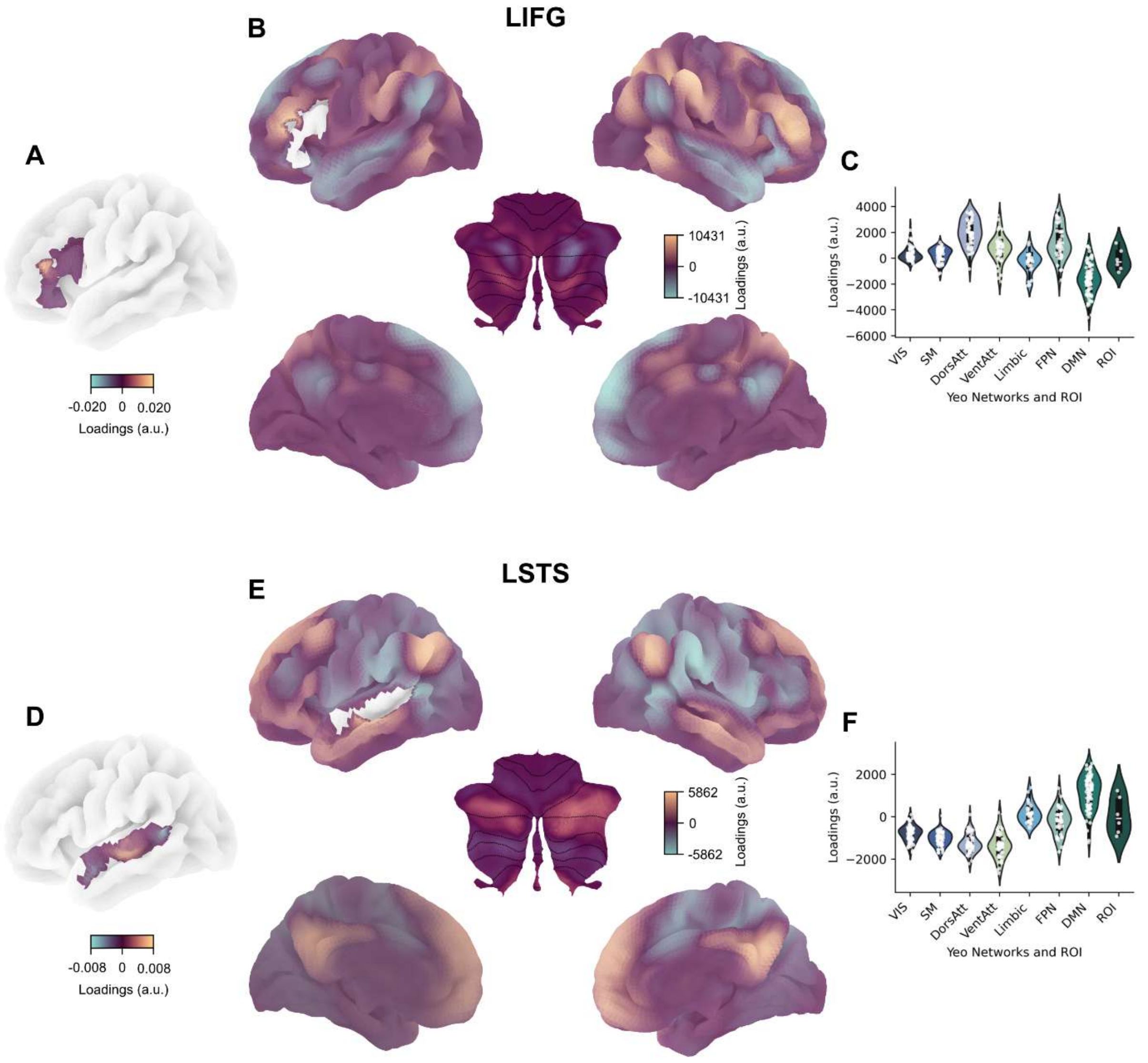


**Supplementary Figure 2. A + D.** Average connectopic map within the ROI (CMAP) for the second gradient. **B + E.** Average connectivity profiles of the ROI to the rest of the brain (PMAP) for the second gradient. **C+F.** Network distributions of the PMAPs based on the Yeo 7 networks and the Glasser atlas for the second gradient. **A-F.** Note that the loadings are arbitrary, but that the sign corresponds between the CMAPs and the PMAPs, meaning that green regions in A+D correspond to green regions in B + E. Abbreviations: LIFG – Left inferior frontal gyrus. LSTS – left superior temporal sulcus. VIS - visual network. SM - somatomotor network. DorsAtt - dorsal attention network. VentAtt - ventral attention network. Limbic - limbic system. FPN - frontoparietal network. DMN - default mode network. ROI - region of interest (corresponding to the LSTS ROI for subplot C and to the LIFG ROI for subplot F). a.u. - arbitrary units.

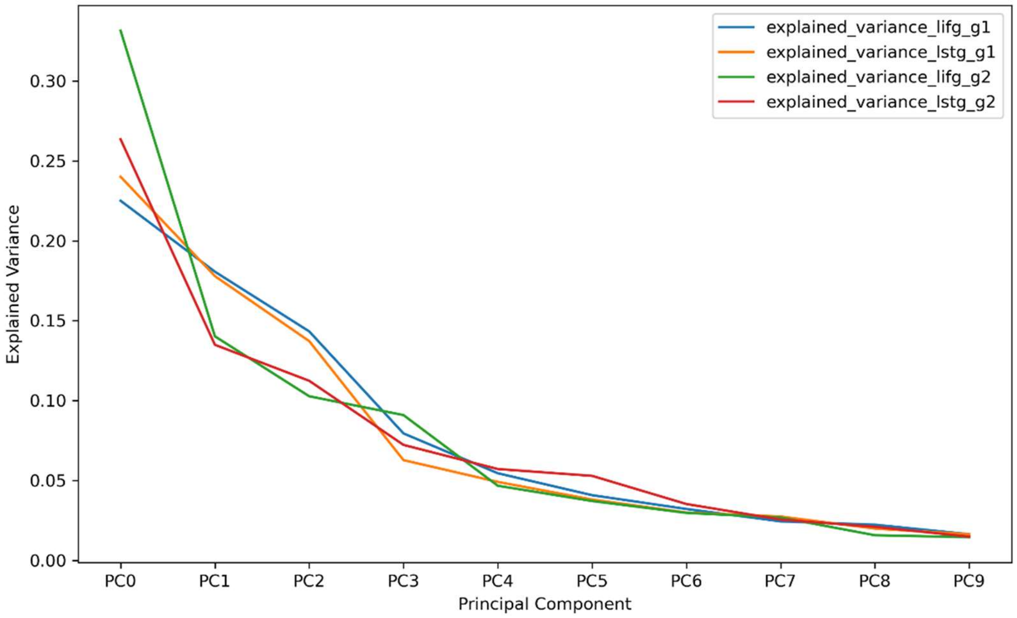


**Supplementary Figure 3.** Scree plots for the principal components of the CMAP and PMAP.

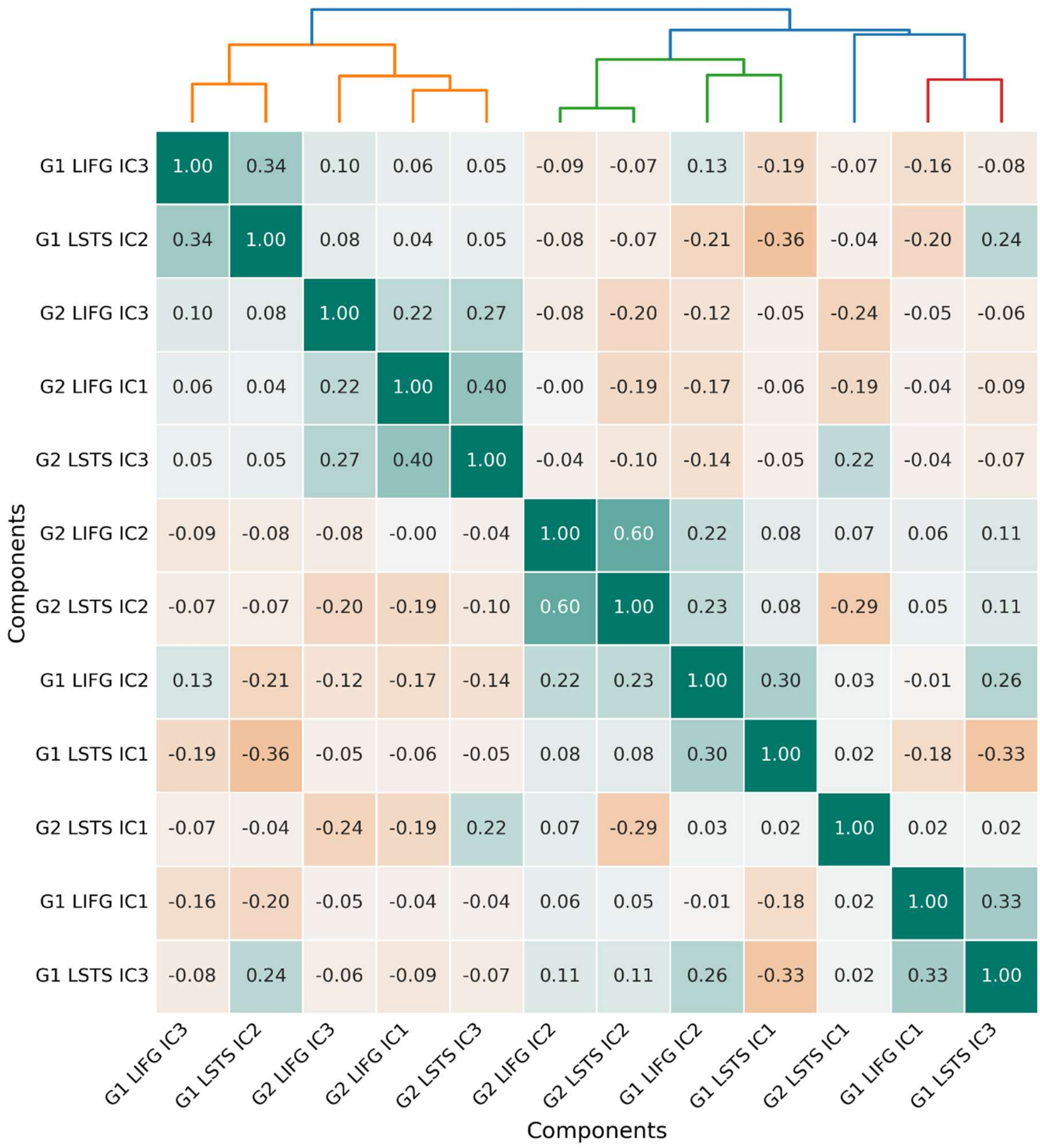


**Supplementary Figure 4.** Hierarchical clustering of components based on subject loading similarities. Hierarchical clustering tree is shown on top of the figure. The clustering can be subdivided in either 2 main clusters or 5 sub-clusters. Both main clusters span both connectopic maps (G1 and G2) and both ROIs (LSTS and LIFG).

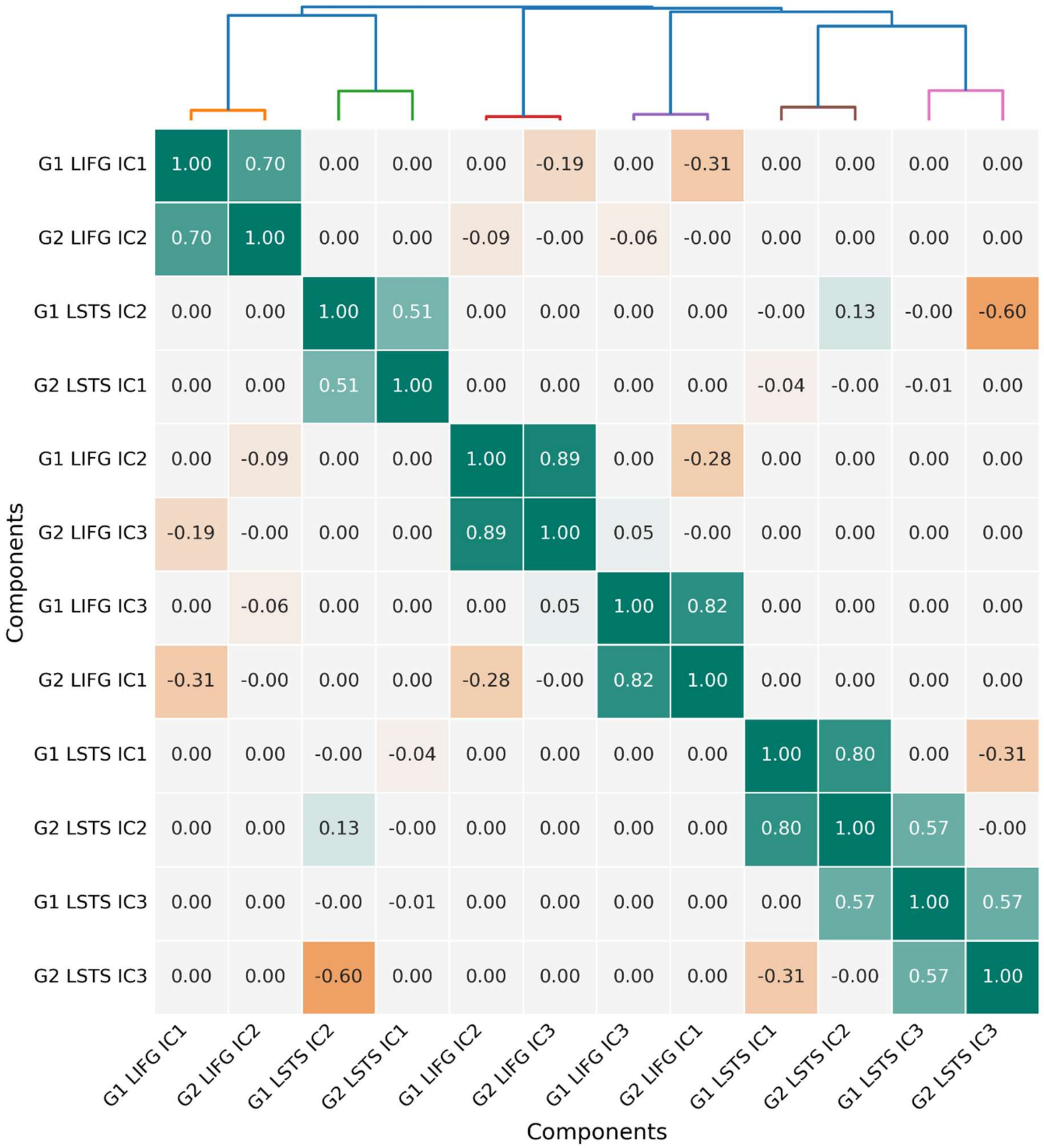


**Supplementary Figure 5.** Hierarchical clustering of components based on spatial similarity of the independent component brain maps. We observe overlap between both connectopic maps (G1 and G2). Note that correlations between both ROIs (LIFG and LSTS) spatial maps are 0 because there are no overlapping voxels. Within-ROI 0 correlations are indicative of the independence criterion of ICA.

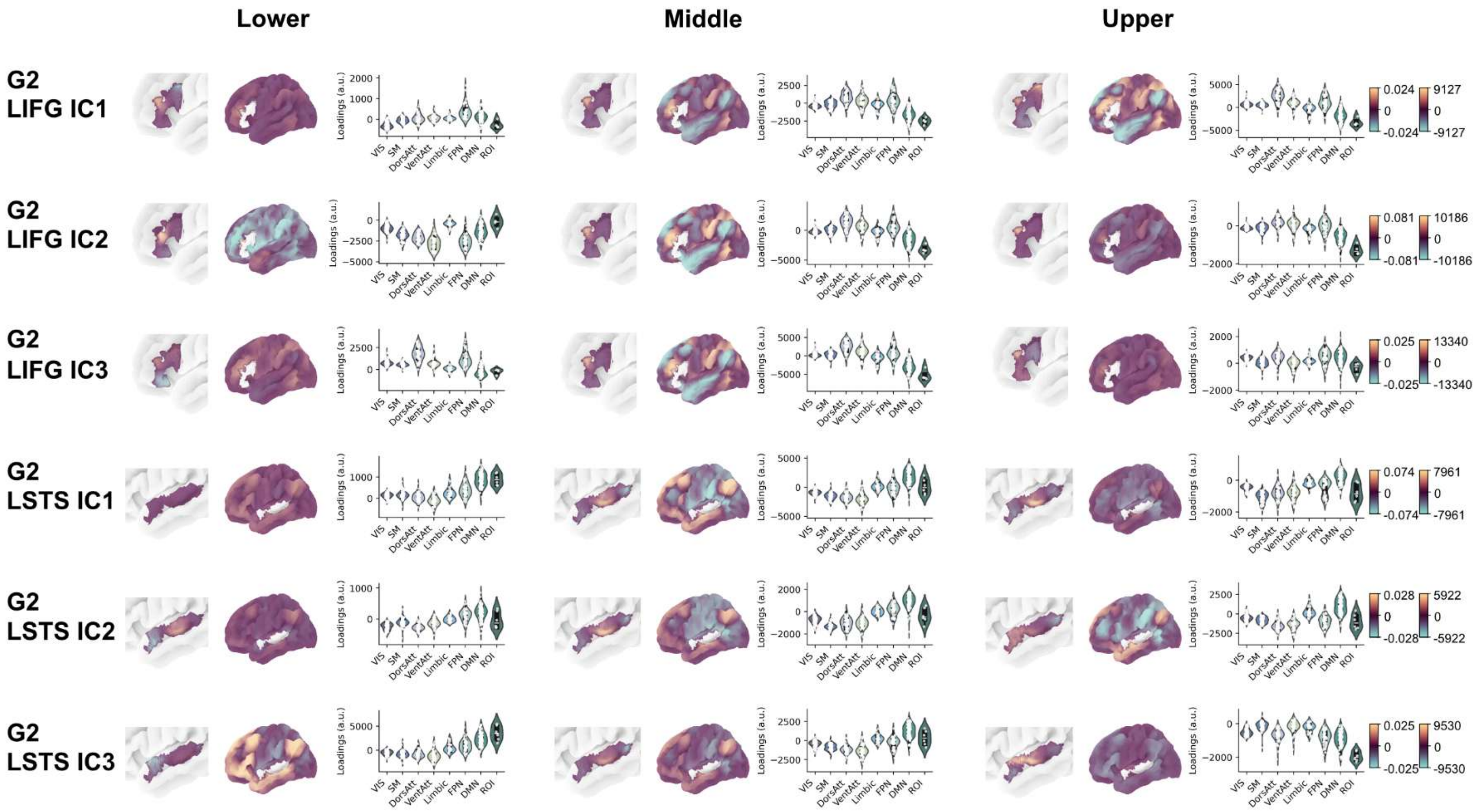


**Supplementary Figure 6.** Population-based variation of CMAP and PMAP differences. The average CMAPs and PMAPs are shown for gradient 2 (G2) for the bottom 50, middle 50 and upper 50 subjects on the IC loadings. Abbreviations: LIFG – Left inferior frontal gyrus. LSTS – left superior temporal sulcus. VIS - visual network. SM - somatomotor network. DorsAtt - dorsal attention network. VentAtt - ventral attention network. Limbic - limbic system. FPN - frontoparietal network. DMN - default mode network. ROI - region of interest (corresponding to the LSTS ROI for subplot C and to the LIFG ROI for subplot F). a.u. - arbitrary units.

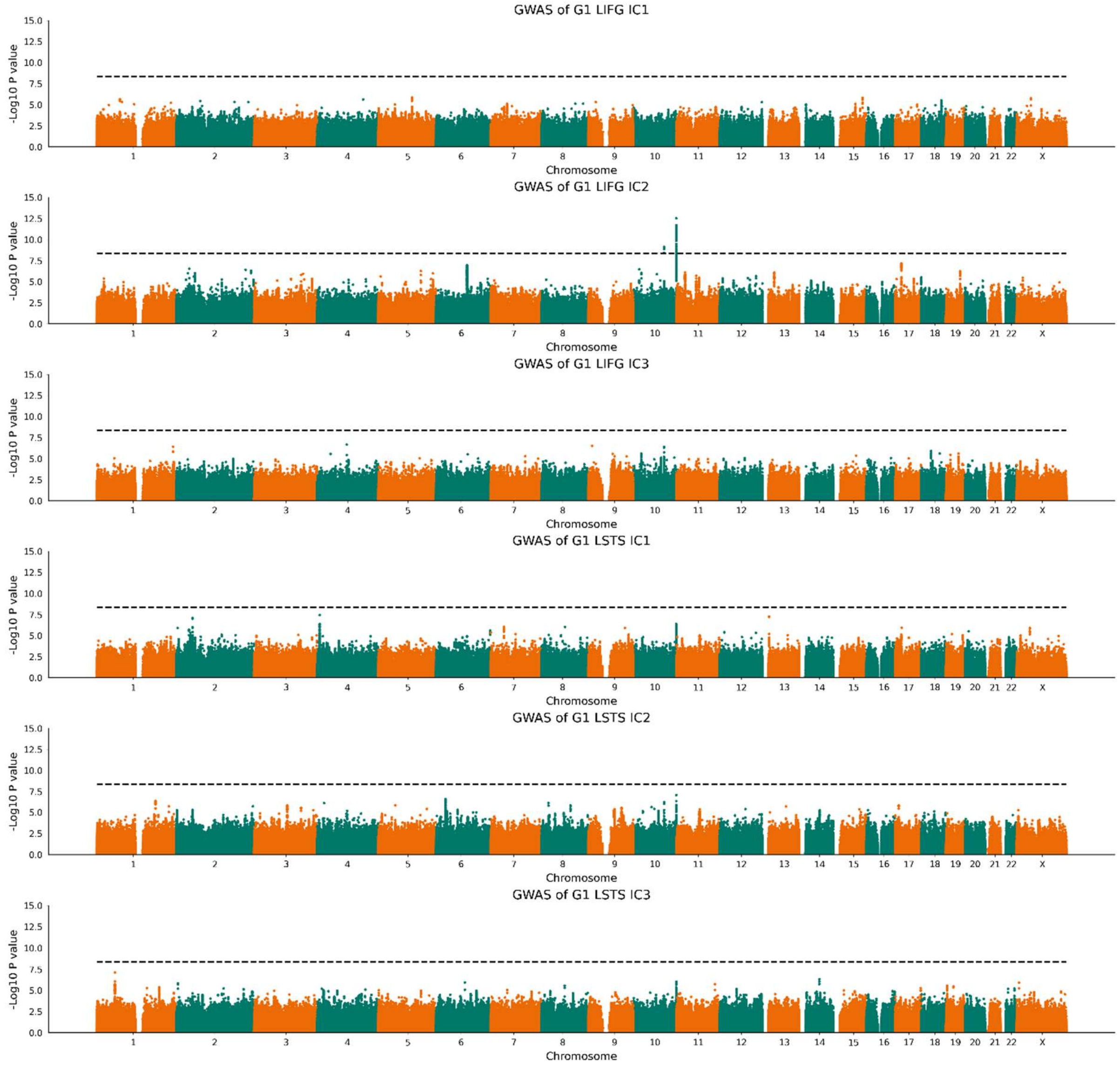


**Supplementary Figure 7.** GWAS plots of all 6 independent components of gradient 1 (N=45,438 for LIFG; N=44,925 for LSTG). X-axis indicates location on the genome, Y-axis indicates association strength by -log10 P-value. Dashed line indicates genome-wide significance corrected for 11 IDPs (P = 4.5e-9).

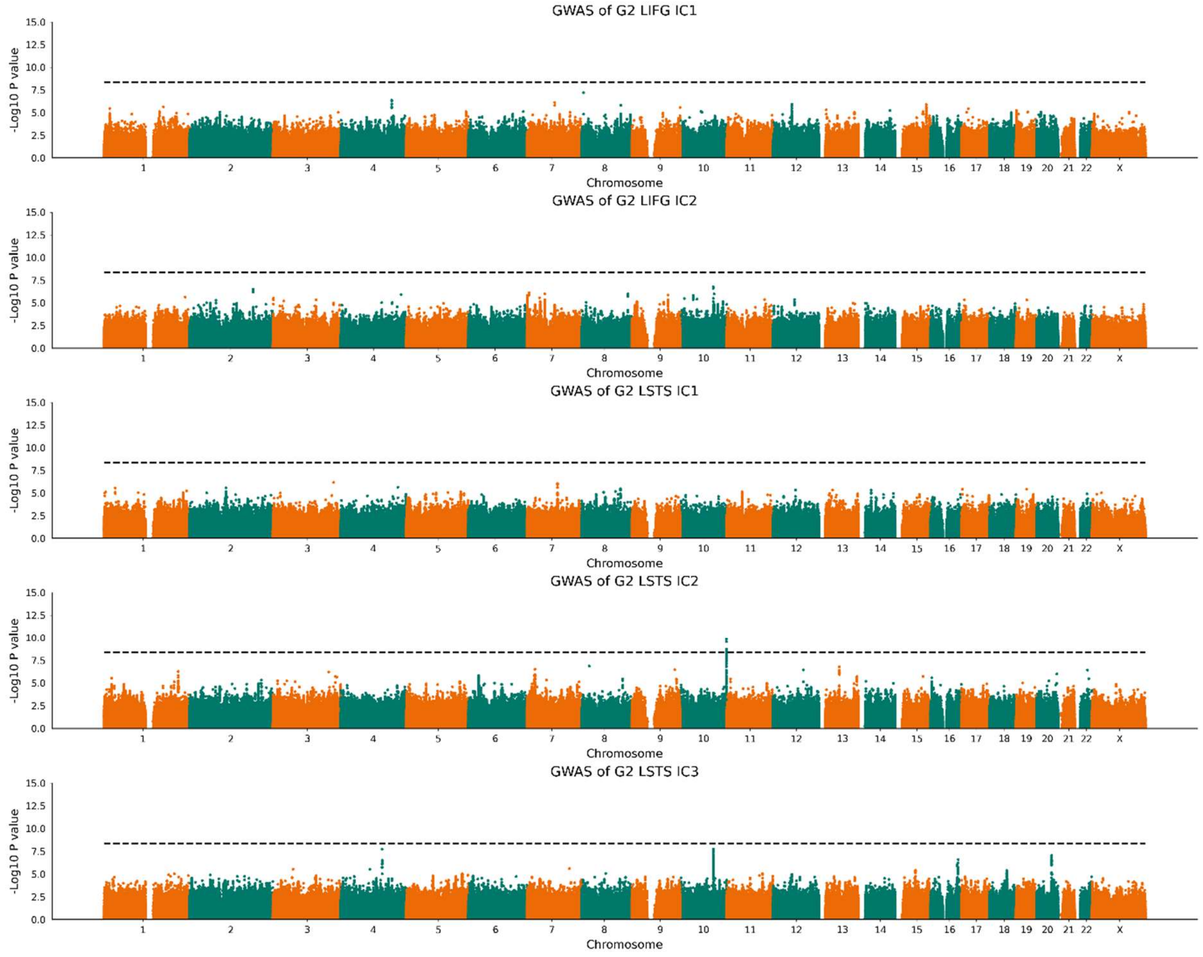


**Supplementary Figure 8.** GWAS plots of all 5 independent components of gradient 2 (N=44,957 for LIFG; N=43,533 for LSTG). X-axis indicates location on the genome, Y-axis indicates association strength by -log10 P-value. Dashed line indicates genome-wide significance corrected for 11 IDPs (P = 4.5e-9).

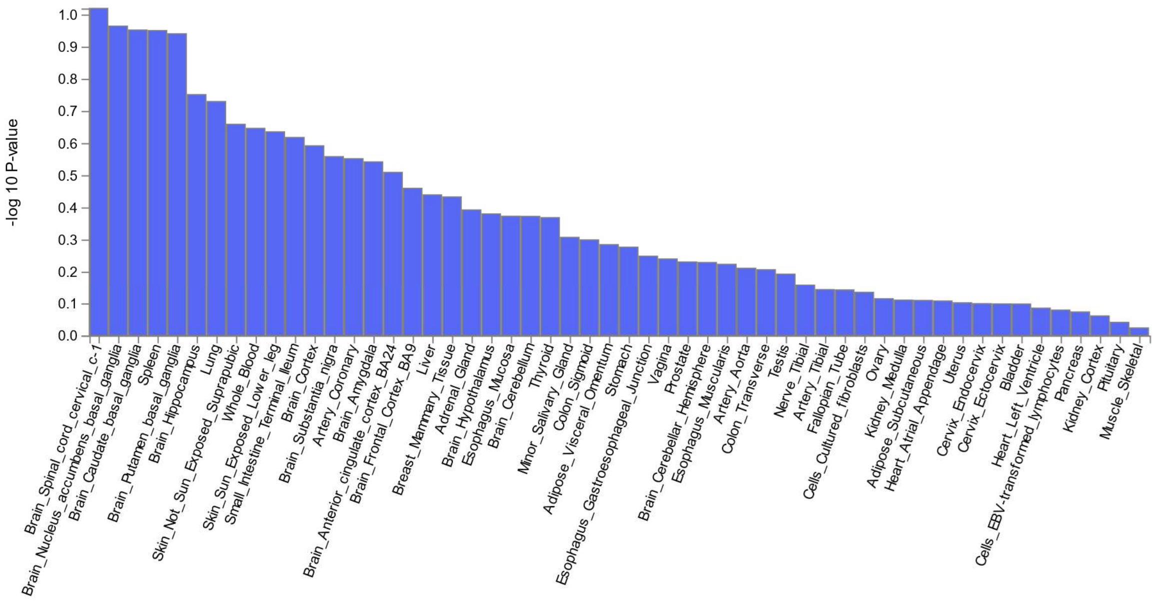


**Supplementary Figure 10.** GTEx results for 51 tissue types based on the mapped genes from the mvGWAS results. Tissue type on the X-axis, association strength (-log10 P-value) on the Y-axis.

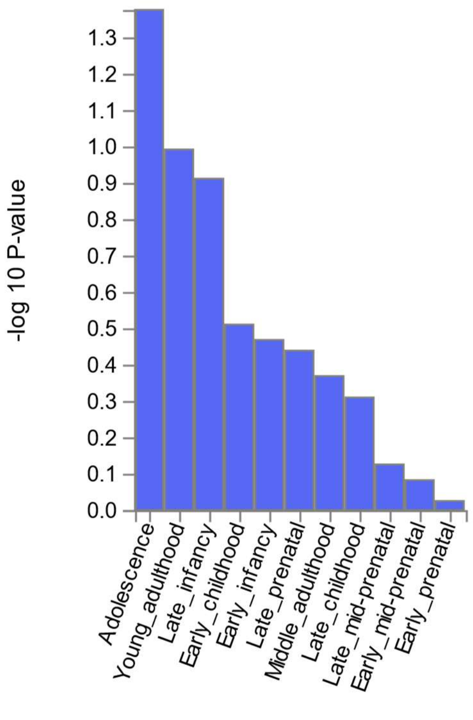


**Supplementary Figure 11.** BrainSpan association results for 11 developmental stages based on the mapped genes from the mvGWAS results. Tissue type on the X-axis, association strength (-log10 P-value) on the Y-axis.

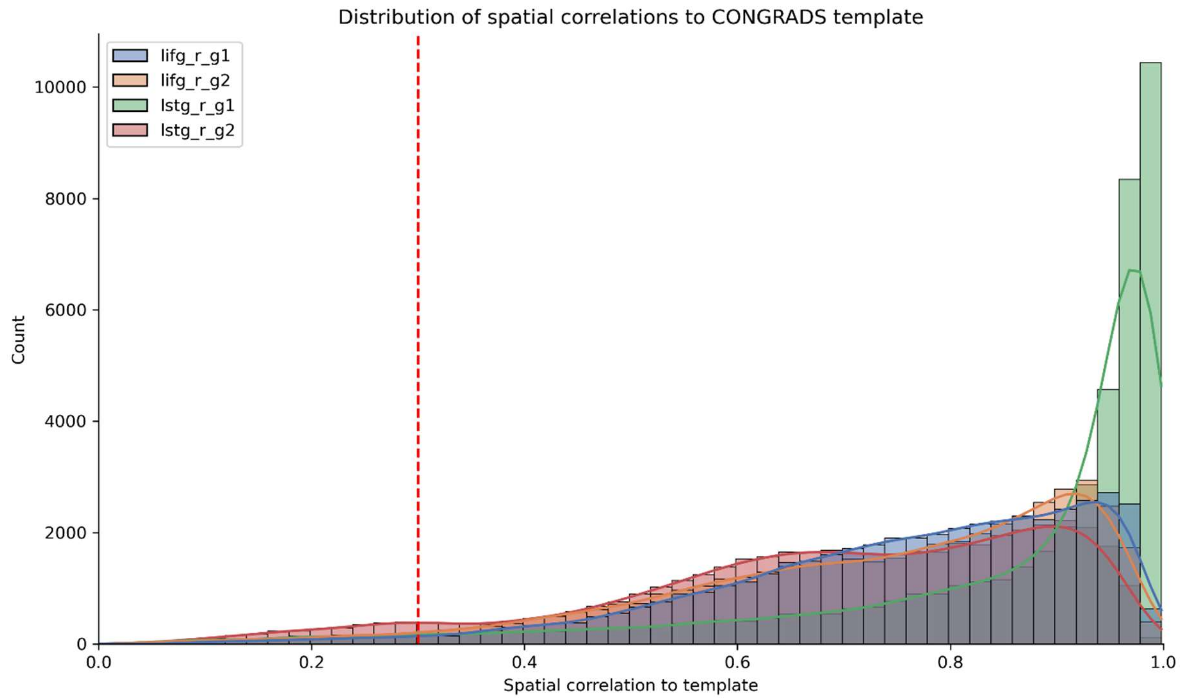


**Supplementary Figure 11.** Distribution of spatial similarity to template across all subjects. Dashed line indicates the threshold for exclusion (r < 0.3).
